# In response to Luteijn *et al*.: Concerns regarding cGAMP uptake assay and evidence that SLC19A1 is not the major cGAMP importer in human PBMCs

**DOI:** 10.1101/798397

**Authors:** Christopher Ritchie, Anthony F. Cordova, Lingyin Li

## Abstract

We previously reported that SLC19A1 is an importer of the immunotransmitter 2’3’-cyclic-GMP-AMP (cGAMP)^1^ by performing a genome wide screen in U937 cells. Soon after, Lutejin *et al*. reported similar findings by conducting a screen in THP-1 cells^2^. While the conclusions of these two studies largely overlap, we arrived at significantly different conclusions regarding how broadly SLC19A1 is used by different cell types. Our study suggests that in addition to SLC19A1, many cultured and primary cell types use alternative, unidentified transporters to import cGAMP and other cyclic dinucleotides (CDNs). This conclusion was based on our findings that inhibition of SLC19A1 did not significantly reduce extracellular cGAMP signaling in multiple cell types, including primary CD14^+^peripheral blood mononuclear cells (PBMCs) from most donors. In contrast, Luteijn *et al*. concluded that SLC19A1 is the major CDN importer in humans, largely based on their use of a radiolabeled [^32^P] cGAMP uptake assay. Using this assay, they showed that inhibition of SLC19A1 abolishes [^32^P] uptake in total PBMCs. However, they did not test whether inhibition of SLC19A1 affects extracellular cGAMP signaling in these cells. Here, we highlight an important issue with the [^32^P] cGAMP uptake assay used by Luteijn *et al*. and demonstrate that measuring extracellular cGAMP signaling through the STING pathway is currently the best method for evaluating cGAMP import. We also show that inhibition of SLC19A1 has no effect on extracellular cGAMP signaling in total PBMCs, confirming that this cell type relies on other transport mechanisms for cGAMP import.

## Results

cGAMP is not stable in serum-containing tissue culture media (hereafter referred to simply as media) due to the presence of its hydrolase ENPP1 in serum^3^, and it is degraded into AMP and P_i_, both of which are reported substrates of SLC19A1^4^. Our analysis using thin layer chromatography (TLC) confirmed that in media a small percentage of [^32^P] cGAMP is degraded into [^32^P] AMP and [^32^P] P_i_ while STF-1623, an ENPP1 inhibitor we developed^5^, efficiently blocks degradation (**Fig. 1a**). Therefore, an uptake assay without ENPP1 inhibitors could be measuring uptake of cGAMP degradation products. This may explain why Luteijn *et al*. were able to measure [^32^P] uptake for up to five hours without reaching equilibrium^2^, despite reports that uptake of other SLC19A1 substrates reaches equilibrium within an hour^4,6^. To investigate further, we found that preincubation of [^32^P] cGAMP in media before performing an uptake assay increased [^32^P] uptake, with a linear correlation between the preincubation time and the amount of [^32^P] uptake. Furthermore, adding STF-1623 to the preincubation abolished the increase in [^32^P] uptake, indicating that the increased [^32^P] uptake is due to degradation of [^32^P] cGAMP (**Fig. 1b**). Importantly, STF-1623 did not inhibit uptake of [^3^H] methotrexate (MTX), a model substrate of SLC19A1, indicating that STF-1623 does not inhibit cGAMP uptake through SLC19A1. Similarly, STF-1084, another ENPP1 inhibitor we developed^5^, only weakly inhibited [^3^H] MTX uptake through SLC19A1 (**Fig. 1c**). In addition, extracellular cGAMP signaling, as measured by phosphorylation of IRF3, is not inhibited by STF-1623 or STF-1084 in U937 cells, which use SLC19A1 as their primary cGAMP importer^1^ (**Fig. 1d**).

**Figure 1.**
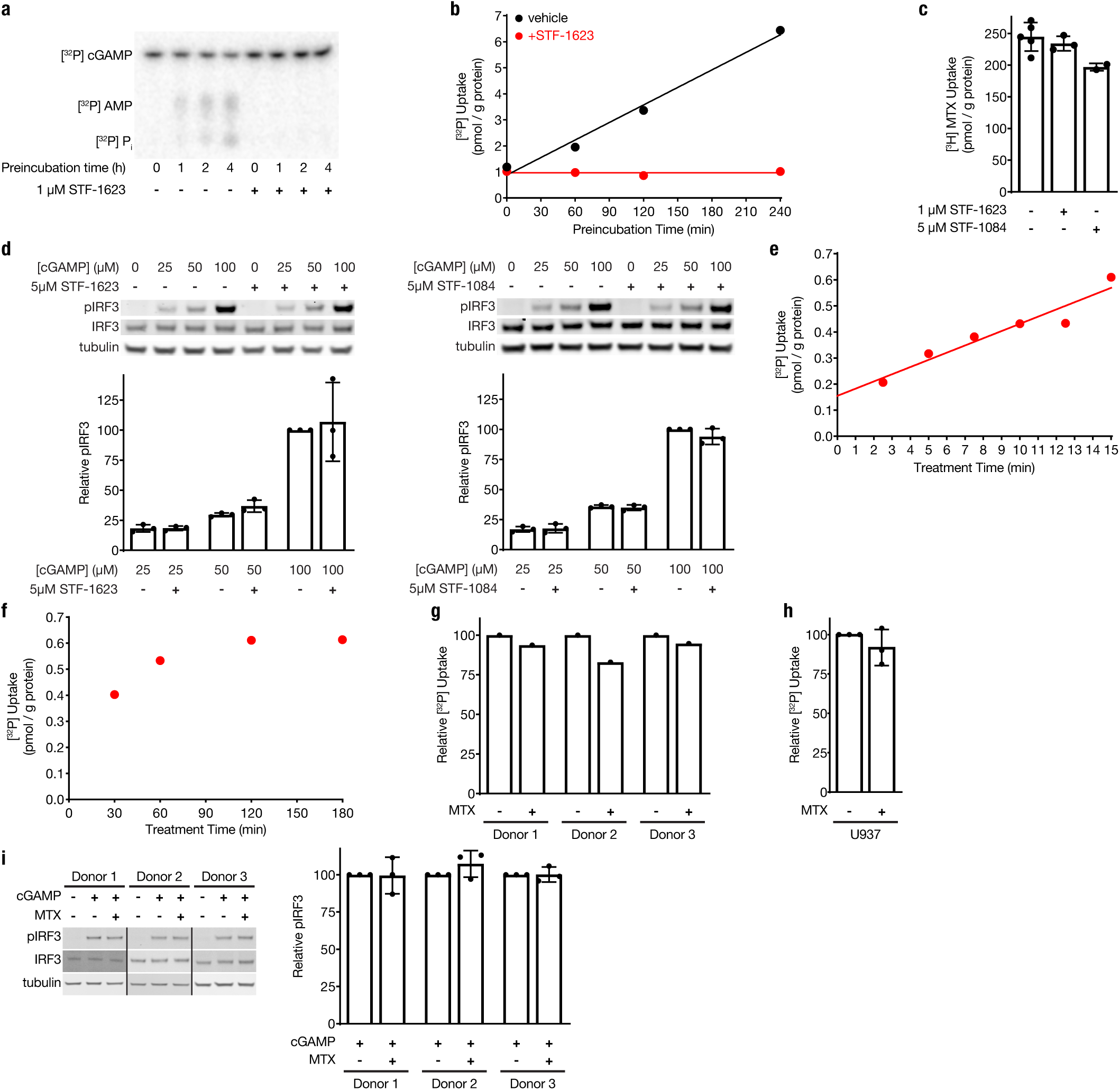
[^32^P] cGAMP uptake assays do not accurately measure cGAMP uptake through SLC19A1. **(a)** TLC analysis of [^32^P] species present after [^32^P] cGAMP preincubation in media. **(b)** Effect of preincubation of [^32^P] cGAMP in media on [^32^P] uptake in U937 cells. 1 nM [^32^P] cGAMP was preincubated for various amounts of time in media in the presence or absence of the ENPP1 inhibitor STF-1623. After this preincubation period, 10^7^ U937 cells were added to the media and [^32^P] uptake was measured after 1 h. **(c)** Effect of ENPP1 inhibitors STF-1623 and STF-1084 on uptake of the SLC19A1 substrate [^3^H] MTX in U937 cells. U937 cells were treated with 17.5 nM [^3^H] MTX for 5 min either alone or in the presence of 1 μM STF-1623 or 5 μM STF-1084 **(d)** Effect of STF-1623 and STF-1084 on extracellular cGAMP signaling in U937 cells. Cells were pretreated with 5 μM STF-1084 or STF-1623 for 15 min, and then treated with the indicated concentrations of cGAMP for 90 min (n = 3 biological replicates for each inhibitor). Data are shown as mean ± SD. **(e)** Short and **(f)** long time courses of 1 nM [^32^P] cGAMP uptake in U937 cells in the presence of 1 μM STF-1623. **(g)** Effect of MTX on [^32^P] cGAMP uptake in PBMCs. PBMCs from three donors were treated with 1 nM [^32^P] cGAMP in the presence or absence of 500 μM MTX. Uptake was measured in the presence of 5 μM STF-1084 for donor 1 and 1 μM STF-1623 for donors 2 and 3. **(h)** Effect of methotrexate on [^32^P] cGAMP uptake in U937 cells. U937 cells were treated with 1 nM [^32^P] cGAMP in the presence or absence of 500 μM MTX. All treatments were performed with 5 μM STF-1084 (n = 3 biological replicates). **(i)** Effect of MTX on extracellular cGAMP signaling in PBMCs. PBMCs from each donor were treated with 50 μM cGAMP in the presence or absence of 500 μM MTX (n = 3 technical replicates). Data are shown as mean ± SD.

Given that our ENPP1 inhibitors effectively block cGAMP degradation without interfering with cGAMP uptake, we used them to evaluate [^32^P] cGAMP uptake kinetics. In the presence of STF-1623, we found that uptake of [^32^P] cGAMP occurred at a linear rate for at least 15 minutes (**Fig. 1e**), with uptake reaching equilibrium by 2 hours (**Fig. 1f**). These results contradict the results presented in Luteijn *et al*., which show linear uptake kinetics for five hours^2^. Taken together, these data strongly suggest that the assay presented by Luteijn *et al*. primarily measured uptake of cGAMP degradation products, rather than cGAMP itself. When we re-evaluated the role of SLC19A1 in PBMCs with our modified [^32^P] cGAMP uptake assay, we found that MTX only weakly inhibits [^32^P] cGAMP uptake in the presence of the ENPP1 inhibitors STF-1084 (donor 1) or STF-1623 (donors 2 and 3) (**Fig. 1g**), in contrast to the 80% reduction observed by Luteijn *et al*.

It is important to note that 1 nM of [^32^P] cGAMP was used in this assay, while we reported that cGAMP has a *K*_i_ of ~700 μM against MTX import by SLC19A1^1^. Therefore, it is possible that that this assay is not operating at a cGAMP concentration that is relevant to cGAMP import by SLC19A1. To test this hypothesis, we performed the same uptake assay using 1 nM of [^32^P] cGAMP in U937 cells. Indeed, [^32^P] cGAMP uptake was not significantly affected by the addition of MTX (**Fig. 1h**). Since most cells require micromolar concentrations of extracellular cGAMP to activate STING signaling, an uptake assay of radioactive cGAMP at nanomolar concentrations is not appropriate to evaluate cGAMP transporters. Therefore, an assay that measures STING pathway activation remains the best available readout for import of extracellular cGAMP. Using this assay, we concluded that SLC19A1 is not the primary cGAMP importer in human CD14^+^ PBMCs^1^ or in total PBMCs (**Fig. 1i**). Additional studies are needed to determine the major cGAMP importer in PBMCs, as well as to identify which cell types use SLC19A1 as their primary cGAMP importer.

## Methods

### Cell Culture

U937 cells were maintained in RPMI (Cellgro) supplemented with 10% heat inactivated FBS (Atlanta Biologicals) and 1% penicillin-streptomycin (GIBCO). Cells were maintained in a 5% CO_2_ incubator at 37 °C.

### Isolation of PBMCs

Buffy coat (Stanford Blood Center) was diluted 1:3 with PBS supplemented with 2 mM EDTA. Diluted buffy coat was layered on top of 50% Percoll (GE Healthcare) containing 140 mM NaCl and centrifuged at 600 × g for 30 min. The separated PBMC layer was collected and washed once with PBS and once with RPMI before being used for uptake and cGAMP signaling experiments.

### Antibodies and reagents

The following primary antibodies were purchased from Cell Signaling Technology and used at a 1:1000 dilution: IRF3 rabbit mAb (cat. #4302), pIRF3 (S396) rabbit mAb (cat. #29047), and α-tubulin mouse mAb (cat. #3873). The following secondary antibodies were purchased from LI-COR Biosciences and used at a 1:15000 dilution: IRDye 680RD Goat anti-Mouse (cat. #926-68070) and IRDye 800CW Goat anti-Rabbit (cat. #925-32211). Radiolabeled [^32^P] ATP (3000 Ci/mmol) used for synthesis of [^32^P] cGAMP was purchased from Perkin Elmer (cat. #BLU003H250UC). Radiolabeled [^3^H] methotrexate was purchased from Moravek (cat. #MT-701). Methotrexate was purchased from Sigma-Aldrich (cat. #A6770). cGAMP was synthesized as previously described^1^. Synthesis of STF-1623 and STF-1084 is described in Carozza *et al*^5^.

### [^32^P] cGAMP synthesis and purification

To catalyze synthesis of [^32^P] cGAMP, recombinant sscGAS was purified as previously described^1^. Synthesis of [^32^P] cGAMP was achieved by incubating the following reaction overnight at room temperature: 50 mM Tris (pH 7.4), 1.67 μM [^32^P] ATP, 100 μM GTP, 20 mM MgCl_2_, 100 μg/ml dsDNA (from herring testes; Sigma-Aldrich), and 50 μM sscGAS. Following synthesis, the reaction was spotted onto a silica TLC plate (Millipore Sigma; cat. # 105548) and run in a solvent containing 85% ethanol and 5 mM NH_4_ HCO_3_ to separate the [^32^P] cGAMP product from the reactants. The band containing the [^32^P] cGAMP product was scraped off the TLC plate and product was eluted from the scraped off silica with water. The eluant was subsequently run through a 3 kDa MWCO filter. To determine the concentration, 10 μL of a 1:100 dilution of the purified [^32^P] cGAMP was added to 10 mL of Bio-Safe II Scintillation Cocktail (Research Products International) to measure [^32^P] activity on an LS 6500 Scintillation Counter (Beckman Coulter). A standard curve was made with the same [^32^P] ATP used in the synthesis reaction to convert [^32^P] activity to molar concentration.

### TLC analysis of [^32^P] cGAMP stability

1 nM [^32^P] cGAMP was incubated in RPMI supplemented with 10% heat inactivated FBS and 1% penicillin-streptomycin in a 5% CO_2_ incubator at 37 °C for the indicated time points. At the end of this incubation period, samples were spotted onto a TLC plate and run in a solvent containing 85% ethanol and 5 mM NH_4_ HCO_3_. Plates were then exposed to a Storage Phosphor Screen (Molecular Dynamics) overnight before being imaged with a Typhoon 9400 Imager (Molecular Dynamics).

### [^32^P] cGAMP uptake assay

10^7^ U937 cells or PBMCs were treated with 1 nM [^32^P] cGAMP in 1 mL of RPMI containing 10% heat inactivated FBS and pen/strep. Short uptake assays (<15 min) were performed in water bath at 37 °C, whereas longer uptake assays (>15 min) were performed in a 5% CO2 humidified incubator at 37 °C. At the end of the treatment time, 10 mL cold PBS were added to cells to halt import. Cells were then pelleted (1000 × g for 5 min), washed twice with 10 mL cold PBS, and lysed with 500 μL 200 mM NaOH. Lysate was completely dissolved by incubating at 65 °C for 45 min. 400 μL dissolved lysate was added to 10 mL of Bio-Safe II Scintillation Cocktail to measure [^32^P] activity. 25 μL dissolved lysate was used for a BCA assay (Thermo Scientific) to measure protein concentration, which was used to normalize [^32^P] activity between samples.

### [^3^H] MTX uptake assay

5 × 10^6^ U937 cells were treated with 17.5 nM [^3^H] MTX in 1 mL of RPMI containing 10% heat inactivated FBS and pen/strep for 5 min in a water bath at 37 °C. At the end of the treatment time, 10 mL cold PBS were added to cells to halt import. Cells were then pelleted (1000 × g for 5 min), washed twice with 10 mL cold PBS, and lysed with 500 μL 200 mM NaOH. Lysate was completely dissolved by incubating at 65 °C for 45 min. 400 μL dissolved lysate was added to 10 mL of Bio-Safe II Scintillation Cocktail to measure [^3^H] activity. 25 μL dissolved lysate was used for a BCA assay (Thermo Scientific) to measure protein concentration, which was used to normalize [^3^H] activity between samples.

### Effect of ENPP1 inhibitors on extracellular cGAMP signaling

5 × 10^5^ U937 cells per sample were pretreated with 5 μM STF-1084 or STF-1623 for 15 min. Cells were then treated with the indicated concentrations of cGAMP for 90 min. Following this, cells were washed once with PBS and then lysed in Laemmli Sample Buffer and run on an SDS-PAGE gel for immunoblot analysis. Densitometric measurements of protein bands were made using ImageJ 1.51g, and tubulin signal was used to normalize pIRF3 signal between samples.

### Effect of MTX on extracellular cGAMP signaling

1 × 10^6^ PBMCs were treated for 2 h with 50 μM cGAMP in the presence or absence of 500 μM MTX. Treatments occurred in 1 mL of RPMI supplemented with 10% heat inactivated FBS and 1% penicillin-streptomycin in a 5% CO_2_ incubator at 37 °C. At the end of the treatment, cells were pelleted, lysed in Laemmli Sample Buffer, and run on an SDS-PAGE gel for immunoblot analysis. Densitometric measurements of protein bands were made using ImageJ 1.51g, and IRF3 signal was used to normalize pIRF3 signal between samples.

## Author Contributions

C.R., A.F.C., and L.L. developed the hypothesis and designed the study. C.R. and A.F.C. conducted the experiments. C.R., A.F.C., and L.L. interpreted and discussed the results. C.R., A.F.C., and L.L. wrote the manuscript.

